# An improved control efficacy against tobacco bacterial wilt by an engineered *Pseudomonas mosselii* expressing the *ripAA* gene from phytopathogenic *Ralstonia solanacearum*

**DOI:** 10.1101/510628

**Authors:** Tao Zhuo, Shiting Chen, Xiaojing Fan, Xun Hu, Huasong Zou

**Affiliations:** Fujian University Key Laboratory for Plant-Microbe Interaction, College of Plant Protection, Fujian Agriculture and Forestry University, Fuzhou, 350002, China

**Keywords:** *Pseudomonas mosselii*, *ripAA*, bacterial wilt, defense signaling pathway, control efficacy

## Abstract

The environmental bacterium *Pseudomonas mosselii* produces antagonistic secondary metabolites with inhibitory effects on multiple plant pathogens, including *Ralstonia solanacearum*, the causal agent of bacterial wilt. In this study, an engineered *P. mosselii* strain was generated to express *R. solanacearum ripAA*, which determines incompatible interactions with tobacco plants. The *ripAA* gene together with its native promoter was integrated into the *P. mosselii* chromosome. The resulting strain showed no difference in antimicrobial activity against *R. solanacearum*. Promoter-LacZ fusion and RT-PCR experiments demonstrated that the *ripAA* gene was transcribed in culture media. Compared with that of the wild type, the engineered strain reduced the disease index by 9.1% for bacterial wilt on tobacco plants. A transcriptome analysis was performed to identify differentially expressed genes in tobacco plants, and the results revealed that ethylene-and jasmonate-dependent defense signaling pathways were induced. These data demonstrated that the engineered *P. mosselii* expressing *ripAA* enables improved biological control against tobacco bacterial wilt by the activation of host defense responses.

**Importance:** Nowadays, the use of biocontrol agents is more and more popular in agriculture, but they cannot replaced of chemical agents mostly, due to the poorer control effect. So the study about how to improve the efficacy of biocontrol agents become necessary and urgent. We increase the efficacy against plant pathogen through introducing an avirulence gene from plant pathogen into the biocontrol agent based on “gene to gene” hypothesis. The new engineered strain can improve the systemic resistance and elicit primary immune response of plants. Our research not only provides a new strategy for genetic modification of biocontrol agent, a number of avirulence gene from pathogen or plant can be tested to be expressed in different biocontrol agents to antagonize plant disease, but also help the study of interaction between phythopathogenic avirulence gene and host.

## Introduction

All major classes of plant pathogens interact with host plants in the “gene-for-gene” model in which a plant resistance (R) protein acts as an elicitor-receptor to directly or indirectly recognize the pathogen-derived *avr* gene product (1). The defense reaction commonly occurs at infection sites to restrict pathogen growth, and in some instances, triggers a hypersensitive response (HR) (2, 3). Upon the recognition of an *avr* gene-harboring pathogen, host cells at the infection site generate an oxidative burst, producing reactive oxygen species, driving cross-linking of cell wall compounds, and leading to the expression of plant genes involved in cellular protection and defense (4, 5). In concert with defense responses, plant R gene-mediated pathogen resistance is used extensively for plant resistance breeding. For example, transduction of the rice *Xa21* resistance gene into banana confers resistance to *Xanthomonas campestris* pv. *musacearum*, which causes the important banana Xanthomonas wilt disease in east and central Africa (6).

Devastating bacterial wilt caused by the *Ralstonia solanacearum* species complex is a major threat to more than 250 plant species in the tropics, subtropics, and other warm temperate areas (7). As all tobacco plant cultivars are suitable hosts for *R. solanacearum* (8), bacterial wilt is one of the most important diseases in tobacco production areas. From 1989 to 1991, in a comprehensive investigation of 16 tobacco production provinces in China, bacterial wilt was found in all provinces except Heilongjiang and Jilin, characterized by high latitudes (9). A high prevalence was found in southern China, especially Fujian, Guangdong, Yunnan, and Guangxi (9). Over time, the disease has become more severe owing to the lack of effective prevention methods.

Like other plant pathogenic bacteria, *R. solanacearum* injects a number of effector proteins into host cells by the type III secretion system (10). Dozens of genomes have indicated a meta-repertoire of effectors in *R. solanacearum* strains (11–13). Among them, the Avr protein RipAA (AvrA) acts as a major host specificity factor recognized by *N. tabacum* and *N. benthamiana* (14, 15). The *ripAA* locus was first cloned from *R. solanacearum* strain AW1 in a 2-kb DNA fragment determining a tobacco incompatibility at the species level (16). Transient expression of RipAA in tobacco leaves induces HR (15). After infiltration into leaves, *R. solanacearum* strains containing the wild-type *ripAA* allele elicit an HR reaction from 16 to 24 h after inoculation, while strains containing a mutated allele cause chlorosis symptoms 36 to 72 h after inoculation, followed by necrosis 48 to 96 h after infiltration (15). The wild-type *ripAA* allele is essential for *R. solanacearum* to induce HR or avirulence in tobacco (14). *R. solanacearum* interacts with tobacco plant species in a “gene-for-gene” relationship depending on wild-type *ripAA*.

As *R. solanacearum* can survive for many years in the environment and invades hosts through plant roots (7), bacterial wilt disease has dramatic effects on contaminated fields owing to limited management measures. In the past few decades, great efforts have been made to reduce disease incidence by the application of antimicrobial agents (17). We previously detected a biological agent with promising control efficacy on tomato bacterial wilt (18). A whole genome sequencing analysis revealed that this agent was a *Pseudomonas mosselii* A1 strain harboring a type III secretion system, like the plant growth-promoting *P. fluorescens* (19). To assess whether the *R. solanacearum* effector RipAA induces resistance to tobacco bacterial wilt, the *ripAA* gene was genetically expressed in *P. mosselii* A1 in this study. Our aim was to construct an engineered *P. mosselii* A1 strain with improved control efficacy in order to facilitate the development of a sustainable protection strategy for tobacco bacterial wilt.

## Materials and methods

### Bacterial and plant materials

The bacterial strains and plasmids used in this study are listed in Table 1. The *P. mosselii* strain was cultivated in Kings B medium at 28°C (20). *R. solanacearum* was isolated from a diseased tobacco plant in Sanmin, Fujian province, China. Nutrient-rich (NB) medium was used to cultivate *R. solanacearum* at 28°C (21). *N. tobacum* NC89 was grown in a greenhouse at 28°C with a photoperiod of 16 h of light (7.5 μmol/m^2^·s) and 8 h of darkness.

**Table 1.** Bacterial strains and plasmids used in this study.

| Strains or plasmids | Relevant characteristics | Resources |
| --- | --- | --- |
| **Strains** |  |  |
| *Pseudomonas mosselii* |  |  |
| A1 | Wild-type *Pseudomonas mosselii* isolated from soil | (18) |
| AA1 | *Pseudomonas mosselii* expressing *RipAA* gene of *Ralstonia solanacearum* | This study |
| A1/pBB:lacZ | Gm<sup>r</sup>, *Pseudomonas mosselii* A1 harboring pBB:lacZ | This study |
| A1/pBB:PlacZ | Gm<sup>r</sup>, *Pseudomonas mosselii* A1 harboring pBB:PacZ | This study |
| *Escherichia coli* |  |  |
| DH5α | *F*'Φ80dlacZDM15D(lacZYA-argF)U169 *endA1 deoR recA1 hsdR17(rK2 mK+) phoA supE44 l2 thi-l gyrA96 relA1* | Clontech |
| *Ralstonia solanacearum* |  |  |
| RsT1 | PB<sup>r</sup>, a *Ralstonia solanacearum* strain isolated from tobacco plants in Sanming Fujian China | Lab collection |
| **plasmids** |  |  |
| pK18mobsacB | Km<sup>r</sup>, suicide vector, *sacB*<sup>+</sup> | (36) |
| pK18:RipAA | Km<sup>r</sup>, a 2.3-kb fusion containing 5' and 3' terminal sequences of a hypothetical gene in *Pseudomonas mosselii* A1, which were interspaced by 1.1-kb DNA fragment of *RipAA* gene with native promoter | This study |
| pBBR1MCS-5 | Gm<sup>r</sup>, 4.7-kb broad-host range plasmid, *lacZ* | (37) |
| pLacZ-Basic | Ap<sup>r</sup>, 7.5-kb pUC replication origin plasmid carrying a β-galactosidase gene | Clontech |
| pBB:lacZ | Gm<sup>r</sup>, pBBR1MCS-5 harboring a 4.6 kb *KpnI/SalI* fragment that contains β-galactosidase gene from pLacZ-Basic | This study |
| pBB:PlacZ | Gm<sup>r</sup>, pBBR1MCS-5 harboring a 276-bp *RipAA* promoter fused with the 4.6 kb *KpnI/SalI* fragment from pLacZ-Basic | This study |

### Integration of *ripAA* into the *P. mosselii* chromosome

Three DNA fragments were cloned in pK19mobSacB to create a construct for integration. The primers used for molecular cloning are listed in Table S1 in the Supplemental Meterial. The first 604-bp of the DAN fragment containing the N terminus of a hypothetical gene was amplified from *P. mosselii* A1 genomic DNA and cloned into pK19mobSacB at *Bam*HI/*Sal*I. An additional *Eco*RI cutting site was introduced in the reverse primer in front of *Sal*I. Second, the 677-bp C-terminal region of the hypothetical gene was cloned into *Eco*RI/*Sal*I. In this PCR amplification, a *Hin*dIII cutting site was introduced in the forward primer behind *Eco*RI. Finally, the DNA fragment containing *ripAA* and its promoter was amplified from *R. solanacearum* GMI1000 genomic DNA and cloned at *Eco*RI/*Hin*dIII. The resulting construct contained the insertion sequence of the hypothetical gene interrupted by the *ripAA* gene. After the introduction of the construct into wild-type *P. mosselii* A1 by electroporation, a standard two-step homologous recombination procedure was used to isolate the marker-free *ripAA* insertion mutant (21).

### Bacterial confrontation assay

The antagonistic activity of *P. mosselii* A1 on tobacco *R. solanacearum* was conducted as described previously (18). *P. mosselii* A1 and tobacco *R. solanacearum* were cultured in Kings B and NB broth medium to OD_600_ = 1.5, respectively. To produce bacterium-containing plates, 1 mL of tobacco *R. solanacearum* cells were added to 100 mL of NA agar medium. After the medium was solidified, 2 μL of *P. mosselii* cells were spotted on each Petri dish and incubated for 3 days at 28°C. The antimicrobial activity of *P. mosselii* was counted based on the inhibitory zone around bacterial colonies, and the diameter of inhibitory zone was recorded using a ruler.

### β-Galactosidase activity driven by the *ripAA* promoter in *P. mosselii* A1

The 276-bp promoter region of *ripAA* was cloned by PCR amplification (Table S1 in the Supplemental Meterial). The fragment was first cloned into the pLacZ-Basic vector (Clontech, Japan) at *Kpn*I/*Xho* sites, and a DNA fragment including the promoter region and *lacZ* gene fusion was digested and moved into pBBR1MCS-5 by digestion with *Kpn*I and *Sal*. Meanwhile, a 4630-bp DNA fragment containing only *lacZ* was generated from empty pLacZ-Basic vector and cloned into pBBR1MCS-5 using the same cutting enzymes *Kpn*I and *Sal*I. The generated pBB:PLacZ and pBB:LacZ constructs were separately introduced into *P. mosselii* A1 by electroporation for β-galactosidase activity analyses according to a standard procedure for the Miller assay (22). For each strain, β-galactosidase activity was analyzed in Kings B or M63 culture conditions.

### Semi-quantitative RT-PCR

Total RNAs were isolated from *P. mosselii* A1 and AA1 cells grown in Kings B or M63 broth using TRIzol reagent (Invitrogen, Carlsbad, CA, USA) according to the manufacturer’s protocol. After contaminated gDNA was removed by treatment with RNase-free DNase (TaKaRa, Otsu, Japan), 2 μg of purified RNA for each sample was used to synthesize cDNA using M-MLV Reverse Transcriptase (Promega, Madison, WI, USA) at 42°C for 1 h. PCR parameters were as follows: 95°C for 5 min followed by 30 cycles of 95°C for 1 min, 60°C for 30 s, and 72°C for 30 s, and a final extension for 10 min at 72°C. The expression of 16S rRNA was determined as a control to evaluate the relative amount of *ripAA* mRNA (Table S1 in the Supplemental Meterial). For each PCR amplification, 8 μL of product was loaded on a 1.5% agarose gel and visualized by ethidium bromide staining.

### Controlling tobacco bacterial wilt using *P. mosselii* AA1

The cultured cells of *R. solanacearum, P. mosselii* A1, and *P. mosselii* AA1 were grown to the mid-logarithmic growth stage at 28°C, harvested at 5,000 × *g* for 10 min, and re-suspended to a final concentration of 1 × 10^8^ CFU/mL. The roots of 4-week-old *N. tobacum* NC89 plants were first wounded and then all inoculated with a *R. solanacearum* cell suspension at 5 mL per pot. Afterwards, tobacco plants were drenched with either *P. mosselii* A1 or AA1. For a negative control, 5 mL of water was applied to the tobacco roots instead of the antagonistic bacterium. All treated plants were grown at 28°C with a soil moisture of > 90%. Disease development was recorded every day until the control plants all completely collapsed. The disease index for each treatment was calculated as described previously (18). Each treatment included 8 plants and was performed in triplicate for statistical analysis.

### High-throughput RNA sequencing

RNA sequencing was performed by GeneBang (Chongqing, China). Root samples were collected from 4-week-old *N. tobacum* NC89 plants at 2 days after inoculation with 5 mL of a 1 × 10^8^ CFU/mL cell suspension of *P. mosselii* A1 or AA1. Total RNAs were extracted from roots using the RNeasy^®^ plant Mini Kit (Qiagen, Shanghai, China) according to the manufacturer’s manual, and mRNAs were purified using poly-T oligo-attached magnetic beads. The concentration, purity, and integrity of RNA were assessed using a NanoDrop™ spectrophotometer (Thermo Scientific, Waltham, MA, USA) and an RNA 6000 Nano Kit with the Bioanalyzer 2100 system (Agilent Technologies, Santa Clara, CA, USA). The TruSeq RNA Library Preparation Kit (Illumina, San Diego, CA, USA) was used to generate RNA libraries, and the quality was assessed using the Agilent Bioanalyzer 2100 system. The plants treated with water were used as a negative control. For each treatment, triplicate experiments were performed for statistical analyses. Thereafter, nine libraries were constructed and were deep sequenced on the Illumina HiSeq 4000 platform. The final reads were aligned to the complete reference genomic of Edwards 2017 in the Solanaceae plants genomics database network (ftp://ftp.solgenomics.net/genomes/Nicotiana_tabacum/edwards_et_al_2017). Genes with log2 fold change ≥ 1 and P ≤ 1 in the comparison between control and treatment groups, which were determined using the DESeq2 (version 1.12.3) package in R (version 3.3.2) and tested for significant differences using Cuffdiff v.2.2.156, were considered differentially expressed. Next, Fisher’s exact test and the false discovery rate (FDR) were used to evaluate each differentially expressed gene (DEG). KEGG analysis was performed using the differentially expressed genes.

### Accession numbers

All the data of transcriptome were deposited with NCBI Sequence Read Archive (SRA) (https://www.ncbi.nlm.nih.gov/sra). The accession numbers of root samples treated by H2O, *P. mosselii* A1 or *P. mosselii* AA1 from three biological repeats were SRR8205398, SRR8205405, SRR8205406, SRR8205399, SRR8205400, SRR8205402, SRR8205401, SRR8205403 and SRR8205404 respectively.

## Results

### Construction of an engineered *P. mosselii* A1 carrying the *ripAA* gene

Owing to the necessity for incompatible interactions with tobacco plants, the *R. solanacearum* effector gene *ripAA* was genetically expressed in *P. mosselii* A1. A hypothetical gene located from 119609 to 120907 in the *P. mosselii* A1 chromosome (GenBank No. NZ_CP024159.1) was chosen as the insertion site to avoid any effects on bacterial growth or antimicrobial activity (Fig. 1A). Three DNA fragments were inserted into the molecular cloning sites of the suicide vector pK19mobSacB, including the left and right parts of the hypothetical gene and a fragment containing the *ripAA* gene with the native promoter. The third fragment was inserted in the middle of left and right parts of the hypothetical gene (Fig. 1B). After the introduction of the resulting recombinant construct into wild-type *P. mosselii* A1, the *ripAA* gene with the native promoter was integrated into the chromosome following two steps of homologous recombination. The engineered *P. mosselii* strain was named AA1 and verified by PCR amplification (Fig. 1C).

**Figure 1.**
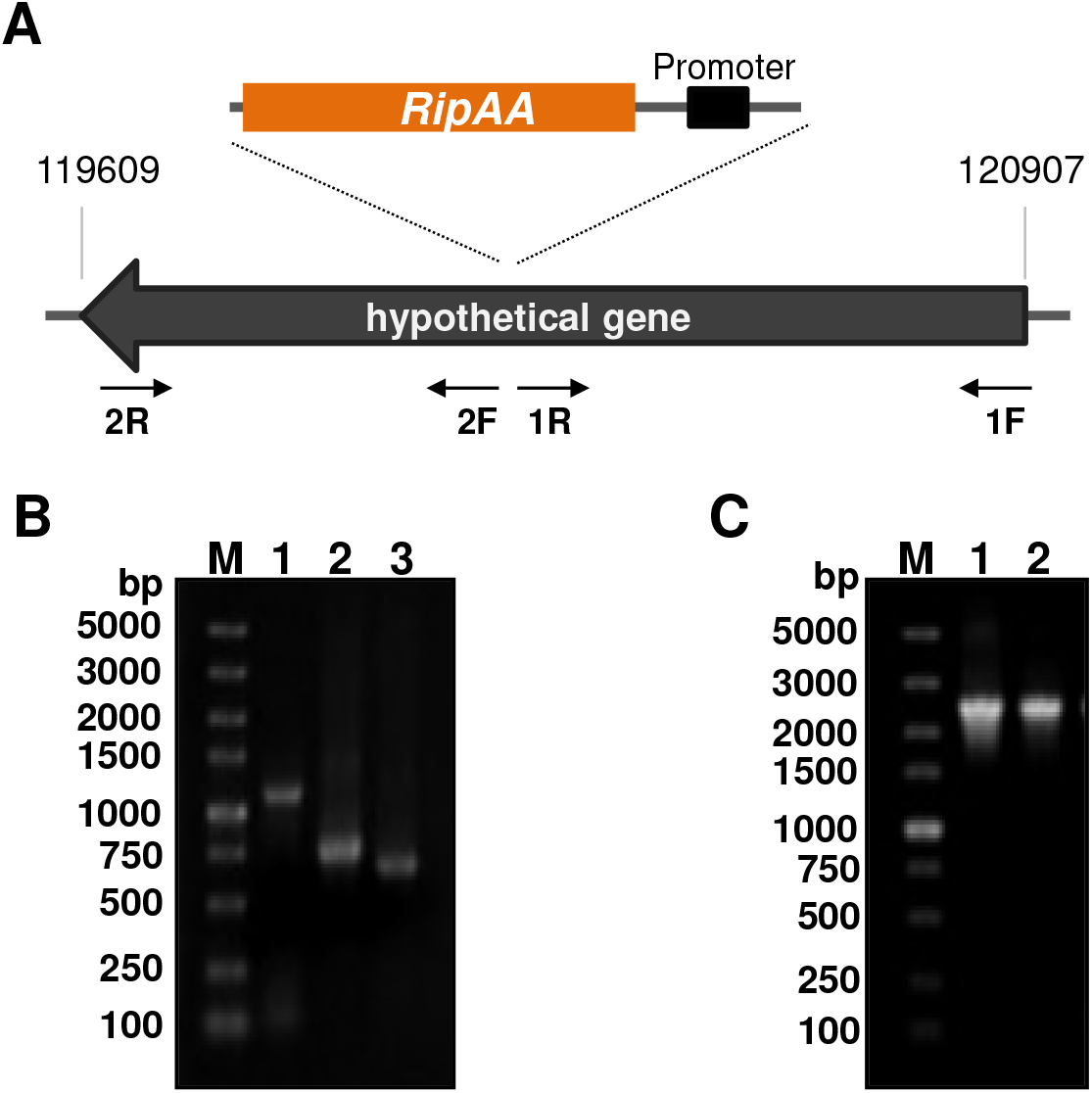
Construction and verification of the engineered *P. mosselii* strain AA1. A. Schematic of the strategy to generate a recombinant AA1 strain carrying *ripAA*. The primer sets used for homologous recombination are indicated by arrows. The insertion site of *ripAA* is shown by dotted lines. B. PCR products cloned in the suicide vector pK19mobSacB. Lane M shows a 5-kb DNA marker, lane 1 shows the PCR product *avrA* from *R. solanacearum*, lanes 2 and 3 show the PCR products obtained using primer sets 1F/1R and 2F/2R from *P. mosselii* A1, respectively. C. PCR verification of engineered *P. mosselii* AA1. Lane M shows a 5-kb DNA marker, lanes 1 and 2 show the PCR products obtained using the primer set 1F/2R from the recombinant vector and *P. mosselii* AA1 genomic DNA. The product in lane 2 was sequenced to confirm that *ripAA* was inserted at the desired location.

### The *avrA* gene is transcribed in *P. mosselii* AA1

To determine whether the *ripAA* gene was transcribed in *P. mosselii* AA1, the promoter activity of *ripAA* was first examined in the *P. mosselii* A1 strain by fusion with *lacZ* gene in the pBBR1MCS-5 vector. In contrast to the negative control harboring the *lacZ* gene without the promoter, the *ripAA* promoter activated *lacZ* gene expression in both Kings B and minimal medium M63. Under either culture medium, yellow colors were clearly observed from the strain carrying the *ripAA* promoter *lacZ* fusion 3 min after the addition of assay buffer containing 2-nitrophenyl-β-d-galactopyranoside (ONPG), and no color was detected from the control strain (Fig. 2A). Quantitative analysis revealed that β-galactosidase activity driven by the *ripAA* promoter was dramatically higher than that of the control (Fig. 2B). This suggested that the *ripAA* promoter was sufficient to drive the transcription of *ripAA* in *P. mosselii* A1. Semi-quantitative RT-PCR was subsequently performed to detect the transcriptional level of *ripAA* in the engineered strain. As shown in Fig. 2c, the transcription of *ripAA* was detected in *P. mosselii* AA1 but not in *P. mosselii* A1 cultured either in Kings B or M63 broth. By contrast, the internal control 16S RNA was expressed with no difference. This demonstrated that *ripAA* was transcribed in *P. mosselii* AA1.

**Figure 2.**
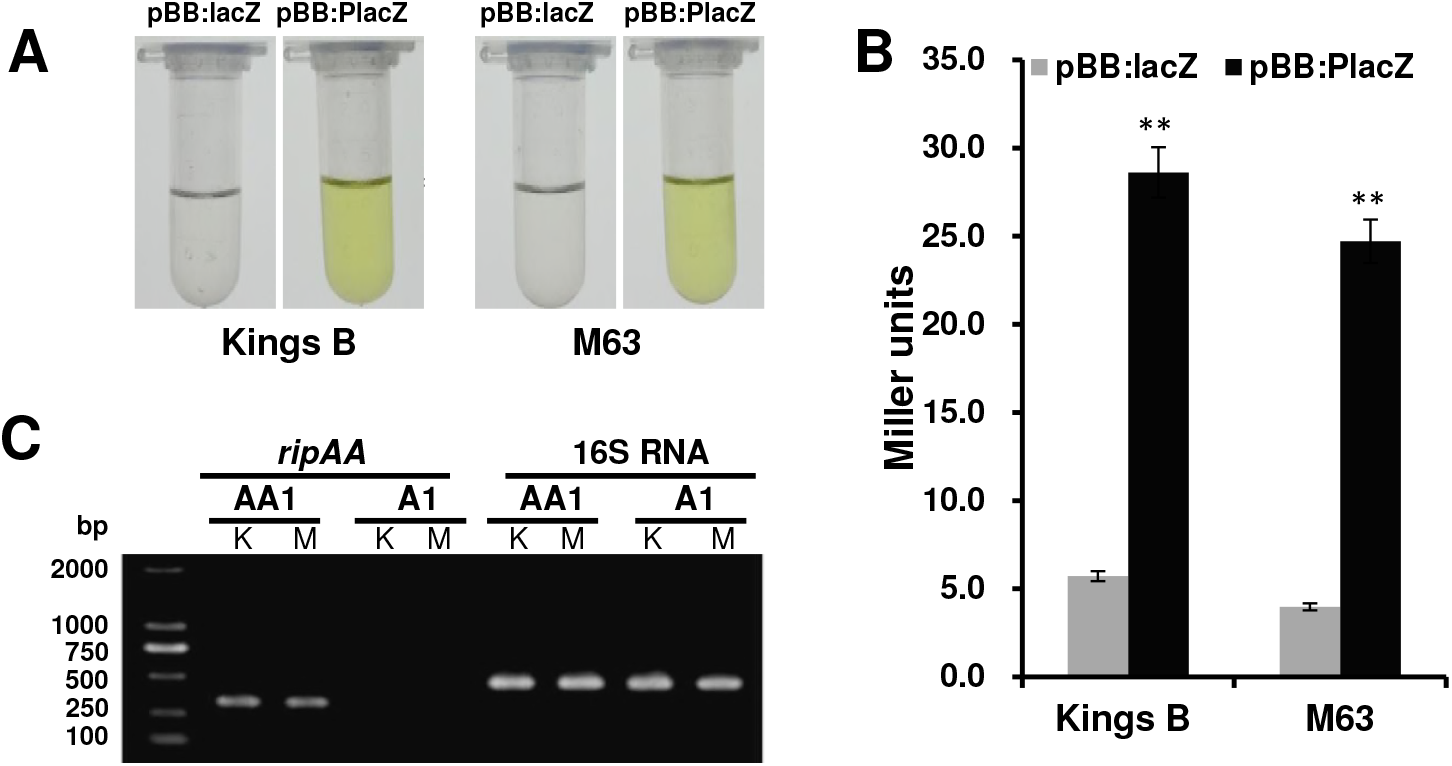
Transcription of *ripAA* in *P. mosselii* AA1. A. A yellow color indicated β-galactosidase activity. The *ripAA* promoter was fused to *lacZ* in the pBBR1MCS-5 vector. The pBBR1MCS-5 vector with only *lacZ* was used as a negative control. After both constructs were introduced into *P. mosselii* A1, β-galactosidase activity was assayed in Kings B or M63 broth. B. Quantitative analysis of β-galactosidase activity driven by the *ripAA* promoter in *P. mosselii* A1. Error bars represent the standard deviation from three independent experiments. Differences were evaluated using Student’s *t*-tests (**P < 0.01.) B. Semi-quantitative RT-PCR analysis of *ripAA* transcription in *P. mosselii* AA1. Total RNAs were isolated from cells grown in Kings B and M63 broth. 16S rRNA was used as an internal control. DNA marker was DL2000 (TAKARA, Dalian, China). K: Kings B medium. M: M63 medium.

### *P. mosselii* AA1 retains antimicrobial activity on *R. solanacearum*

A bacterial confrontation assay was conducted on plates to evaluate whether the insertion of *ripAA* had an effect on antimicrobial activity or not. Like the wild-type *P. mosselii* A1, the engineered *P. mosselii* AA1 expressing *ripAA* showed antimicrobial inhibition activity on the growth of *R. solanacearum*. Inhibition zones were clearly observed around the colonies of both *P. mosselii* AA1 and A1 at 3 days post-inoculation (Fig. 3A). The diameters of the inhibition zones were approximately 1.5 cm for both strains (Fig. 3B). A statistical analysis indicated no difference between *P*. *mosselii* A1 and AA1, indicating that the expression of *ripAA* did not affect the antimicrobial activity of *P. mosselii* AA1.

**Figure 3.**
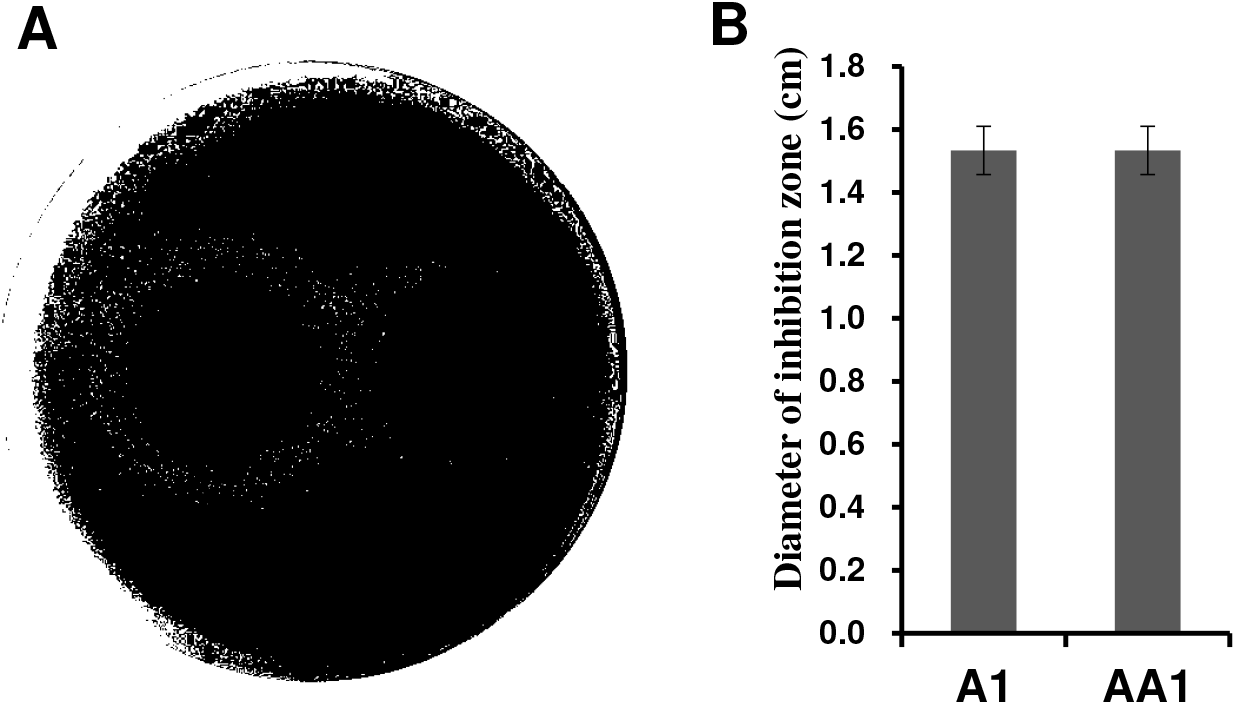
Antimicrobial activity of *P. mosselii* A1 and its mutant against *Ralstonia solanacearum* in a plate confrontation experiment. A. Plate confrontation on *Ralstonia solanacearum* by *P. mosselii* AA1. The cultured *P. mosselii* cells were prepared to OD_600_ =1.5, and 2 μg of cell suspension was spotted on NA plates containing *R. solanacearum*. The inhibitory effect was recorded at 3 days postinoculation. B. Diameters of inhibitory zones, as determined at 3 days postinoculation from three replications. Error bars represent the standard deviation from three independent experiments.

### *P. mosselii* AA1 increases control efficacy on tobacco bacterial wilt

*P. mosselii* AA1 was applied to root-wounded tobacco plants to assess control efficacy. In the negative control (water treatment), *R. solanacearum* resulted in wilt disease symptoms on tobacco plants at 3 days post-inoculation, whereas the plants receiving A1 and AA1 treatment showed wilt symptoms at 4 and 5 days post-inoculation, respectively (Fig. 4). The plant disease index for the water treatment control increased sharply to 92.9 at 15 days post-inoculation (Fig. 4). The disease index for A1 treatment decreased to 38.5 and that of the AA1 treatment decreased to 29.4. In contrast to the A1 treatment, the plants receiving AA1 treatment exhibited a delay of 1 day for wilt symptom appearance; the disease index was reduced 9.1% at 15 days post-inoculation (Fig. 4).

**Figure 4.**
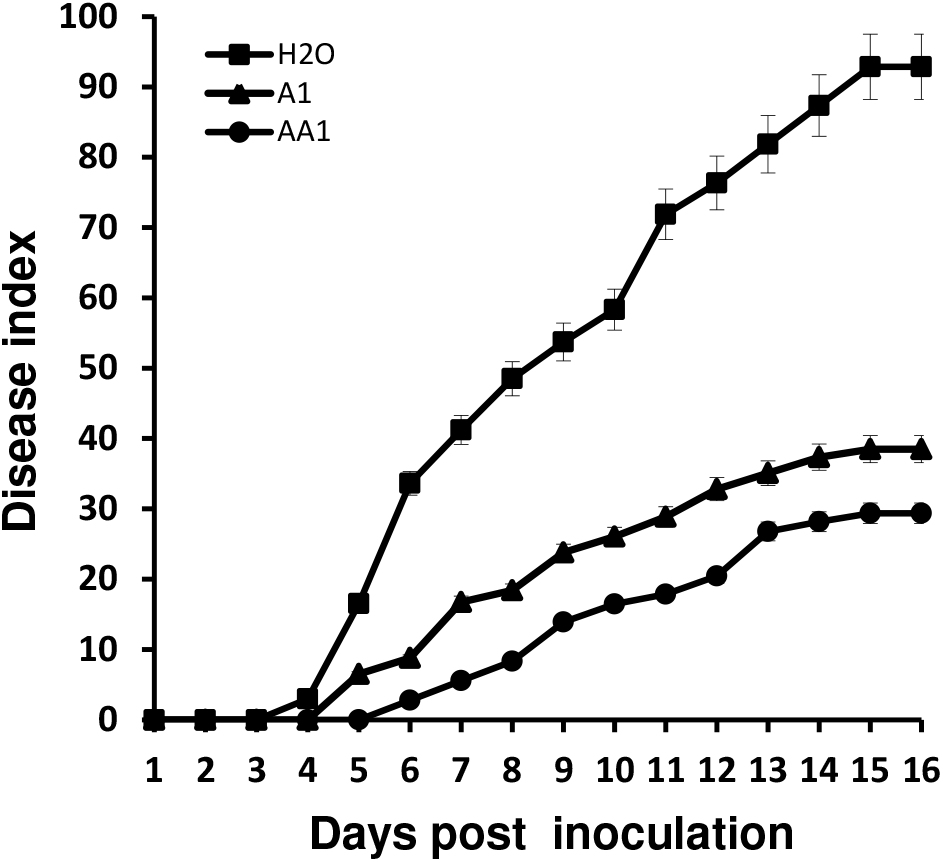
Progression of bacterial wilt on tobacco plants treated with AA1. The disease rate was recorded daily from 0 to 16 days after wounded root inoculation. Each point represents the mean disease rate of 8 inoculated plants per treatment. Error bars represent the standard deviation from three independent experiments.

### Transcriptome analysis of tobacco plants treated with *P. mosselii* AA1

DeSeq2 was used to identify tobacco genes that were differentially expressed in tobacco plants treated with the AA1 strain. The transcription data for plants receiving AA1 treatment were analyzed by comparisons with both H2O and A1 treatments. In the comparison with the H2O treatment, 2497 genes were differentially expressed, including 1296 up-regulated and 1201 down-regulated genes (Fig. 5A). In the comparison with the A1 treatment, 1277 genes were up-regulated and 1112 genes were down-regulated. The genes that showed the same expression changes for the H2O and A1 treatments were recognized as potential genes affected by *ripAA* (Fig. 5A). In total, 890 genes were up-regulated and 635 were down-regulated in plants receiving AA1 treatment in comparisons with both A1 and H2O (Fig. 5A). These 1525 differentially expressed genes were proposed to be specifically affected by *ripAA*.

**Figure 5.**
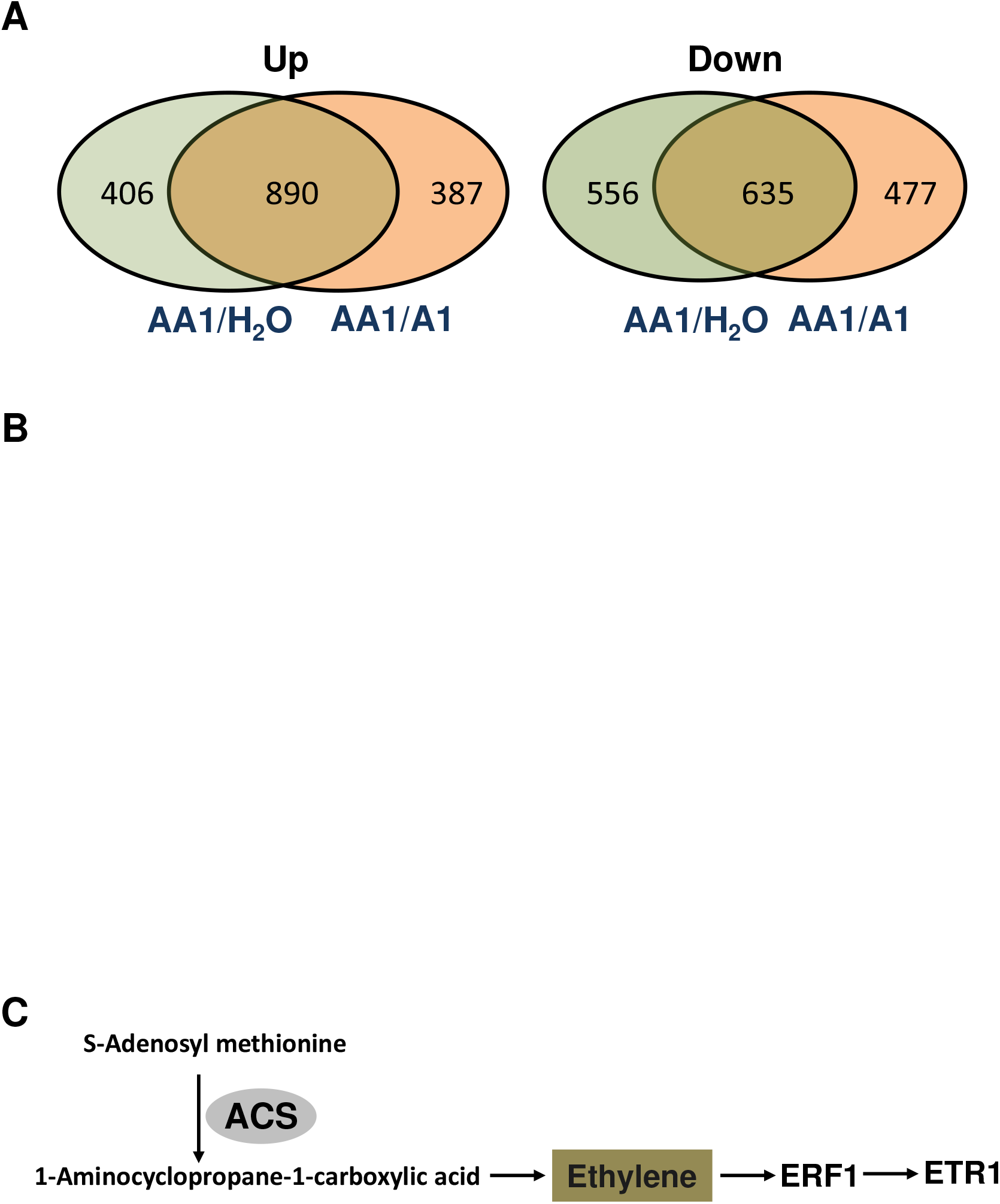
Identification of differentially expressed genes by RNA sequencing. A. Diagram showing the numbers of genes that shared the same patterns of transcriptional change. Genes with log2 fold change ≥ 1 and P ≤ 1 for the comparison between control and treated groups were determined using the DESeq2 package in R (version 3.3.2) and Cuffdiff v.2.2.156. B. KEGG classification of differentially expressed genes. C. Induced genes involved in ethylene signaling pathway.

Among the 1525 unknown differentially expressed genes, 256 up-regulated and 54 down-regulated genes were classified into 31 functional pathways by a KEGG pathway enrichment analysis (Fig. 5B). Among the 310 enriched genes, 79 genes were classified into several functional groups, implying the versatile biological functions of the genes. Furthermore, 89 genes were enriched in the metabolic pathway, which had the largest number of genes among 31 enriched pathways (Fig. 5B). Four enriched pathways were up-regulated by *ripAA*, including starch and sucrose metabolism, phenylpropanoid biosynthesis, pyrimidine metabolism, and MAPK signaling pathway-plant pathways (Fig. 5B). Alanine, aspartate and glutamate metabolism, peroxisome, carotenoid biosynthesis, and glycosphingolipid biosynthesis pathways were down-regulated.

The ethylene and jasmonate signaling pathways were activated by AA1 expressing *ripAA*. The up-regulation of 1-aminocyclopropane-1-carboxylate synthase (*ACS*) (Nitab4.5_0002381g0080.1) and down-regulation of methyl jasmonate esterase (*MJAE*) (Nitab4.5_0000382g0070.1) suggested that a metabolic increase in ethylene biosynthesis occurred in tobacco plants receiving AA1 treatment. Accordingly, downstream signaling transduction was activated, coupled by the elevated expression of *efr1* and *etr1* (Fig. 5C). For the JA signaling pathway, seven jasmonate ZIM domain-containing genes were down-regulated.

## Discussion

Numerous integrated measures have been established to control bacterial wilt (17) because the pathogen *R. solanacearum* can persist for many years in soil, water, weeds, and diseased plant tissues (7). To avoid chemical contamination in the soil, biological control methods have been rapidly developed in recent years (23–26). *Pseudomonas* spp., *Bacillus* spp., *Streptomyces* spp., and other bacterial species are promising for controlling bacterial wilt (17). Interestingly, several avirulent *R. solanacearum* isolates are able to prevent bacterial wilt disease development (27). *Pseudomonas* spp. are optimal biocontrol agents as they compete for root colonization, synthesize allelochemicals, and induce systemic resistance of host plants (28). We recently reported a biocontrol agent, *Pseudomonas* spp. strain A1, with antimicrobial activity against *R. solanacearum* (18). Despite a remarkable reduction in disease development, the disease index for plants receiving *Pseudomonas* spp. A1 treatment still remains about 40 at 20 days after inoculation, suggesting that an improved strategy is needed (18).

*Pseudomonas* spp. A1 was initially assigned to the *P. putida* group based on a phylogenetic analysis of the 16S rRNA sequence (18). Based on the complete genome sequence completed in our lab, it was definitively identified as *P. mosselii*. Since the initial discovery from clinical specimens in 2002 (29), *P. mosselii* isolates have been found extensively in soil and water environments (30, 31). In addition to the abilities to degrade a wide variety of aromatic chemicals, *P. mosselii* strains show excellent antagonistic activity against plant pathogens (32). Based on the complete genome sequence information, *P. mosselii* A1 studied here possesses the type III secretion system, which is widely distributed in plant-colonizing bacteria, including *P. fluorescens* and *P. putida* (19). In *P. fluorescens* SBW25, the constitutive expression of *hrpL* and *avrB* confers the ability to induce HR in *A. thaliana* ecotype Col-0 (19). In this study, hetero-expression of the *R. solanacearum ripAA* gene in *P. mosselii* A1 resulted a constitutive expression module and thus improved the control efficacy against tobacco bacterial wilt.

*R. solanacearum* RipAA acts as an avirulent factor, together with RipP1, to determine the resistance reaction of tobacco to pathogens according to the “gene-for-gene” model (15). By expressing *ripAA* in *P. mosselii* A1, ethylene and jasmonate signaling pathways in tobacco plants were significantly induced relative to levels in plants treated with *P. mosselii* A1. A previous study has reported that the knockout of Haem peroxidase enables *P. putida* to induce systemic resistance (33). As Haem peroxidase is also found in *P. mosselii* A1, we proposed that the wild-type A1 strain possesses the ability to induce systemic resistance. Furthermore, a total of 12 Haem peroxidase genes in tobacco plants were up-regulated in response to the expression of *ripAA*, suggesting that oxidative reactions are activated in tobacco plants. The expression of *ripAA* in *P. mosselii* A1 induced several defense-related signaling pathways to improve tobacco resistance to bacterial wilt caused by *R. solanacearum*.

In *R. solanacearum*, the transcription of *ripAA* is under the control of the key regulator HrpB, which is responsible for the initiation of *hrp* genes, required for the type III secretion system assembly (34). The expression of *hrpB* is induced in plants or in an hrp-inducing, nutrient-poor medium (35). Even though the transcription of *ripAA* requires induction by environmental signals (15), the *ripAA* promoter drove *ripAA* transcription in *P. mosselii* A1 cultured in either Kings B or M63 medium. The transcription of *ripAA* in the engineered AA1 strain was detectable by a sensitive RT-PCR analysis. This suggested that the *hrpB* homolog (CSH50_RS09230) in *P. mosselii* A1 was sufficient to activate the transcription of *ripAA*. Unfortunately, we failed to detect the epitope-tagged RipAA-Myc in the AA1 strain and thus could not determine whether it was delivered to the extracellular milieu. We speculated that the RipAA protein was not present in sufficient quantities for a western bot analysis. Irrespective of this, the engineered AA1 strain expressing *ripAA* improved plant resistance to bacterial wilt.

Our original aim was to create an engineered strain possessing both antimicrobial activity and defense induction abilities to prevent bacterial wilt disease at diverse infection stages. At the invasion stage, *P. mosselii* AA1 inhibited *R. solanacearum* in the rhizosphere; it also promoted plant defense to the pathogen after colonization in the vascular system. Even though the disease index decreased by about 9.1%, the engineered AA1 did not fully prevent disease occurrence. There are two explanations for this observation. First, the insertion of *ripAA* in the engineered AA1 strain did not enhance antimicrobial activity against *R. solanacearum*, which could not lead to a decreased pathogen population in the soil. Second, the incompatible interaction of *R. solanacearum* with tobacco plants is determined by both RipAA and RipP1 (15). The expression of only the *ripAA* gene in *P. mosselii* AA1 could not induce sufficient resistance to bacterial wilt in tobacco plants.

In conclusion, we constructed an engineered *P. mosselii* strain AA1 expressing the *ripAA* gene from *R. solanacearum*, which is involved in incompatible interactions with tobacco plants. *P. mosselii* AA1 exhibited improved biological control efficacy against tobacco bacterial wilt by evoking several defense signaling pathways. These results not provide insight into defense mechanisms induced by *ripAA*, but also provide a basis for improving the management of bacterial wilt disease using pathogen-derived Avr proteins.

## Acknowledgements

This study was funded by the National Science Foundation of China (31671988 and 31701752) and the Scientific Project of Fujian Provincial Education Department (JA15152).

